# Machine Learning exploratory technic detected that men might be up to eight times more affected by the control effect and three times more affected by the placebo effect than women

**DOI:** 10.1101/2023.04.02.535269

**Authors:** Roberto Lohn Nahon, José Kawazoe Lazzoli, Marcio Vinícius de Abreu Verli, Luis Carlos Oliveira Gonçalves, Aníbal Monteiro de Magalhães Neto

## Abstract

**Background:** The Placebo effect has been historically described since the beginning of Medicine. When the most skeptical researchers say they do not believe in Noetic Science but use a placebo in their research, they generate an apparent contradiction. The present study aimed to understand the noetic influence on high-level athletes, using a sportomics strategy, statistical exploratory techniques of machine learning and holistic analysis.

**Methods:** The study included 14 volunteer volleyball athletes. Each volunteer was submitted to four running tests of 3,000 meters, on a 400-meter track, with one test each subsequent day. On the first day, the athletes performed the first test of 3,000, aiming to adapt to the trial (ADAPT 1), and on the second day, the same adaptation (ADAPT 2). On the third and fourth days, the placebos were introduced, and on the third day, the athletes received the information that that would be just a placebo, which was called (CONTROL). On the fourth day, when the identical placebo was given, the athletes received the information that it would be a new cutting-edge nutritional supplement being studied (PLACEBO).

**Results:** Men might be up to eight times more affected by the control effect and three times more by the placebo effect than women. Regarding performance, there was an antagonistic behavior concerning gender for the control effect and an agonistic effect for the placebo effect, but with less impact on women. Men also showed a faster adaptation to the test.

**Conclusion:** Noetic science, always considered but never assumed by researchers, is confirmed when the present study reveals that men are more affected by the control effect and the placebo effect than women, with antagonistic behavior concerning gender for the control effect and an agonist effect for the placebo effect, but with less impact on women about performance.

## Introduction

The Placebo effect (PE) has been historically described since the beginning of Medicine (Shapiro, 2007). Its best-known effect is that associated with the use of drugs for some kind of clinical treatment. PE is so important that the standard procedure for investigating the effect of some clinical intervention involves a control group using placebo (an inert substance) or another standard drug for the condition in discussion (Simon, 2000), ideally with a double-blind design (neither the evaluator nor the evaluated person knows whether the substance is or not the tested drug), preventing possible biases to influence the result, a fact that could occur if there was only the “intervention group”, without the control placebo.

Such an effect is hardly quantified biochemically, or by other complementary exams, however it is easily identified clinically, when assessing pain improvement, quality of life improvement or improvement in daily activities. Many of the treatments whose gains are not observed in the perspective of scientific evidence, has its effects associated with PE.

Noetic Science investigates metaphysical phenomena related to the subjectivity of human consciousness (Pekala, 2015). The evolutionary levels of consciousness vary from affective forms to the most advanced cognitive forms of neural processing, ranging from anoetic (knowledgeless) consciousness based on affective feelings to noetic (knowledge-based) and autonoetic (higher reflective mental) processes that allow for perception. conscious (Vandekerckhove & Panksepp, 2011).

One of the most effective applications of this science is in the health area, where studies already point to its contribution as complementary medicine through the insertion of holistic practices, providing comprehensive care and thus ensuring greater efficiency in the therapeutic process (Sawni & Breuner, 2017). In addition, although it is still a science on the rise, it can correlate its object of study to significant advances in several areas of knowledge (Pekala, 2016).

The studies using the sportomics strategy aim to reproduce the natural conditions of sports or training methods for an integrated evaluation to answer different questions, from the immunometabolic stress caused the effect of supplements, among others (Gonçalves et al., 2012; Gonçalves et al., 2022).

The present study aimed to understand the noetic influence on high-level athletes, using a sportomics strategy, statistical exploratory techniques of machine learning and holistic analysis.

## Methods

### Subjects

The study included 14 volunteer volleyball athletes, ten females and four males, with more than 10,000 hours of sports competitions, duly registered in their sports federations and considered high-level athletes in their clubs. Each volunteer was submitted to a pre-participation risk questionnaire and evaluated regarding the use of any medication or treatment for any disease. Exclusion criteria were the use of any medication, any illness or trauma, or the desire not to perform the tests at maximum intensity.

### Experimental Design

Each volunteer was submitted to four running tests of 3,000 meters, on a 400-meter track, with one test each subsequent day. As these tests would be applied aiming at the internal classification in their teams, motivation remained high in all tests.

The distance of 3,000 meters was chosen based on the observation that maximum aerobic power tests should ideally last 8 to 12 minutes, now widely used in modern exercise tests (Buchfuhrer, 1983).

On the first day, the athletes performed the first test of 3,000, aiming to adapt to the trial (ADAPT 1), and on the second day, the same adaptation (ADAPT 2).

On the third and fourth days, the placebos were introduced, and on the third day, the athletes received the information that that would be just a placebo, which was called (CONTROL).

On the fourth day, when the identical placebo was given, the athletes received the information that it would be a new cutting-edge nutritional supplement being studied (PLACEBO).

The time of each athlete in the execution of the cited test in each experimental moment (ADAPT 1, ADAPT 2, CONTROL, PLACEBO) was recorded.

### Statistical Analysis

Initially, descriptive statistics were performed on the data, with measurements of position (mean, median, mode, and percentiles) and dispersion (amplitude, variance, standard deviation, and standard error).

Afterward, the univariate analysis of these data was performed using the Shapiro-Wilk normality test (because the sample was smaller than 30 individuals). The equal variance test would be applied if the Shapiro-Wilk test presented a result indicating normal distribution (P>0.05). For results with

P>0.05, the paired T-Student test would follow; if P≤0.05, the paired T-Student test would follow the non-parametric Mann-Witney test. If the Shapiro-Wilk test presented a result indicating non-normal distribution (P≤0.05), the non-parametric Mann-Witney test would be applied directly.

Still, in the phase of the univariate analysis, the analysis of repeated measures ANOVA One Way dependent was performed because they were the same individuals in different conditions and moments.

At the end of the univariate analysis phase, it became clear that individuals (male and female) could not be evaluated together to investigate the placebo effect (Figure 1).

**Figure 1.**
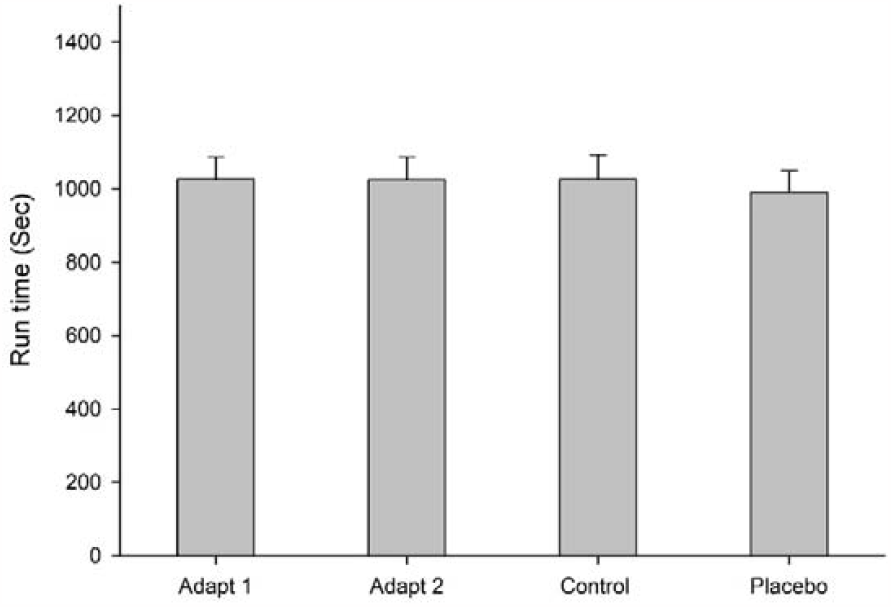
When the effect is observed in a single group, there is no difference in yield (T test; Anova; Cohen d and r).

So, for a better interpretation of the data, the individuals were divided into two groups according to their sex. Then, the calculation of percentage variation was applied:

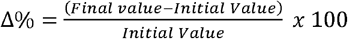

Cohen’s equations (Cohen, 1992) were used to calculate the effect size for all variables to obtain Cohen d and r values:

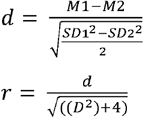

Where, M represent the means of observations and SD their respective standard deviations.

**Table 1.** Values of effect size.

| Effect size | Small | Medium | Large |
| --- | --- | --- | --- |
| **Cohen r** | 0.10 | 0.30 | 0.50 |
| **Cohen d** | 0.20 | 0.50 | 0.80 |

Source: Cohen, 1992.

Next, multivariate data analysis was performed using data mining and machine learning techniques.

In this phase, in order to seek a bivariate measure between the data, because the observations contain quantitative values, the Pearson and Spearman correlation tests were applied, with the Spearman correlation being used for a visual analysis using the heat map strategy and the Pearson test as an initial measure for the following machine learning analyses.

As exploratory models of machine learning: CLUSTER - Classical Clustering (Agglomerative Hierarchical Method) and Nearest neighbor (single linkage); ORDINATION – Principal component Analysis (PCA) and Correspondence Analysis (CA).

The Z score was not previously applied because the observations contained similar measurement units.

SigmaPlot 14.5 (Academic Perpetual License - Single User – ESD Systat® USA) and, Past 4.03 (Free version for Windows) were used to carry out the different statistical tests and produce the graphs.

### Ethical Approval

This study was conducted according to the guidelines laid down in the Declaration of Helsinki, and the experiment met the requirements of research using human subjects (National Health Council, 2012). This study was approved by the Ethics and Research Committee (number 2,230,073) of the Federal University of Mato Grosso (UFMT) and was registered at clinicaltrials.gov (NCT 03522883). The individuals were informed that they could withdraw from the study at any time. Written informed consent was obtained from each subject, who was instructed on the nature of the research and the procedures involved.

## Results

As expected, when they were treated together (men and women), no difference was found for the control or placebo effect (Fig 1). This fact led to the creation of two groups by sex.

After such separation, it became clear that men might be up to eight times more affected by the control effect (Fig 2A) and three times more affected by the placebo effect (Fig 2B) than women.

**Figure 2.**
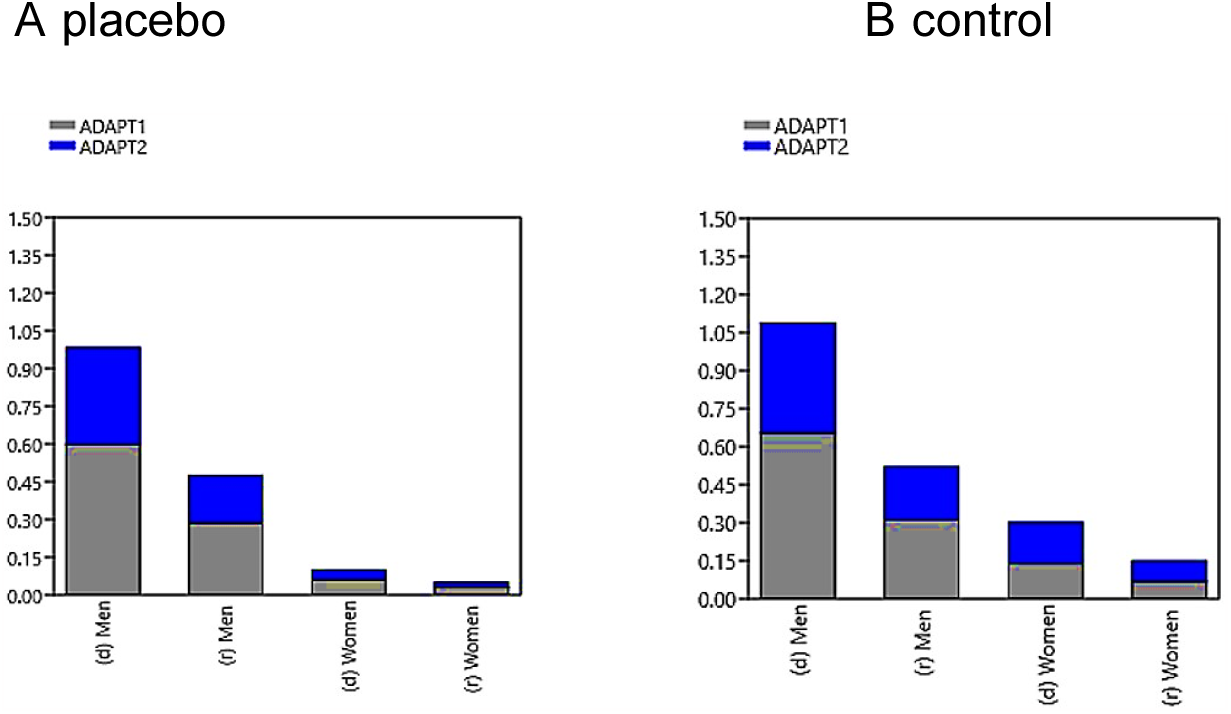
Men might be up to eight times more affected by the control effect (Fig 2A) and three times more affected by the placebo effect (Fig 2B) than women.

Regarding performance, there was an antagonistic behavior concerning gender for the control effect and an agonistic effect for the placebo effect, but with less impact on women. Men also showed a faster adaptation to the test (Fig 3).

**Figure 3.**
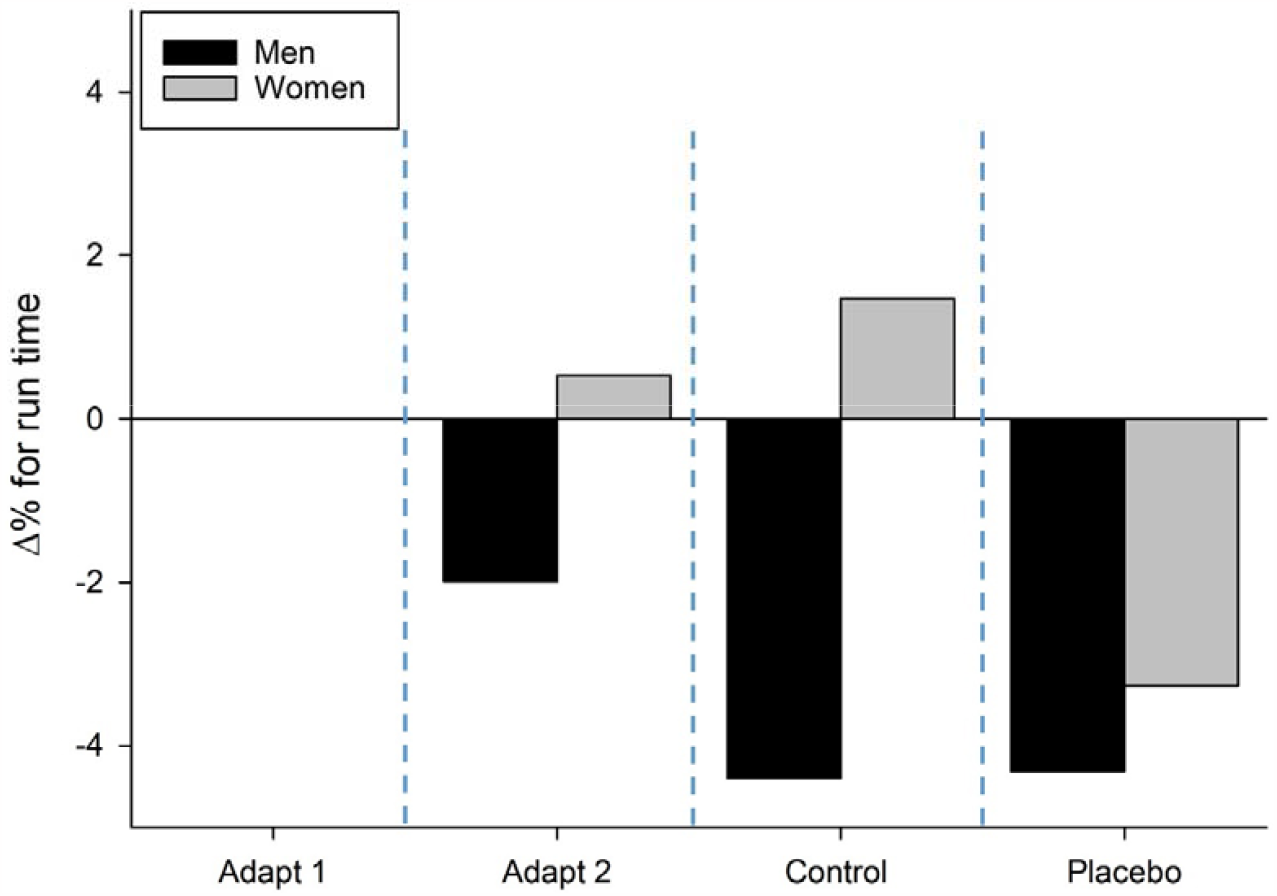
Regarding performance, there was an antagonistic behavior concerning gender for the control effect and an agonistic effect for the placebo effect, but with less impact on women. Men also showed a faster adaptation to the test.

Spearman’s rank correlation test confirmed the choice of separating men from women (Fig 4), because the heatmap revealed two squares of behavior similar to each other and different between the groups of men and women, with high correlations within each group.

**Figure 4.**
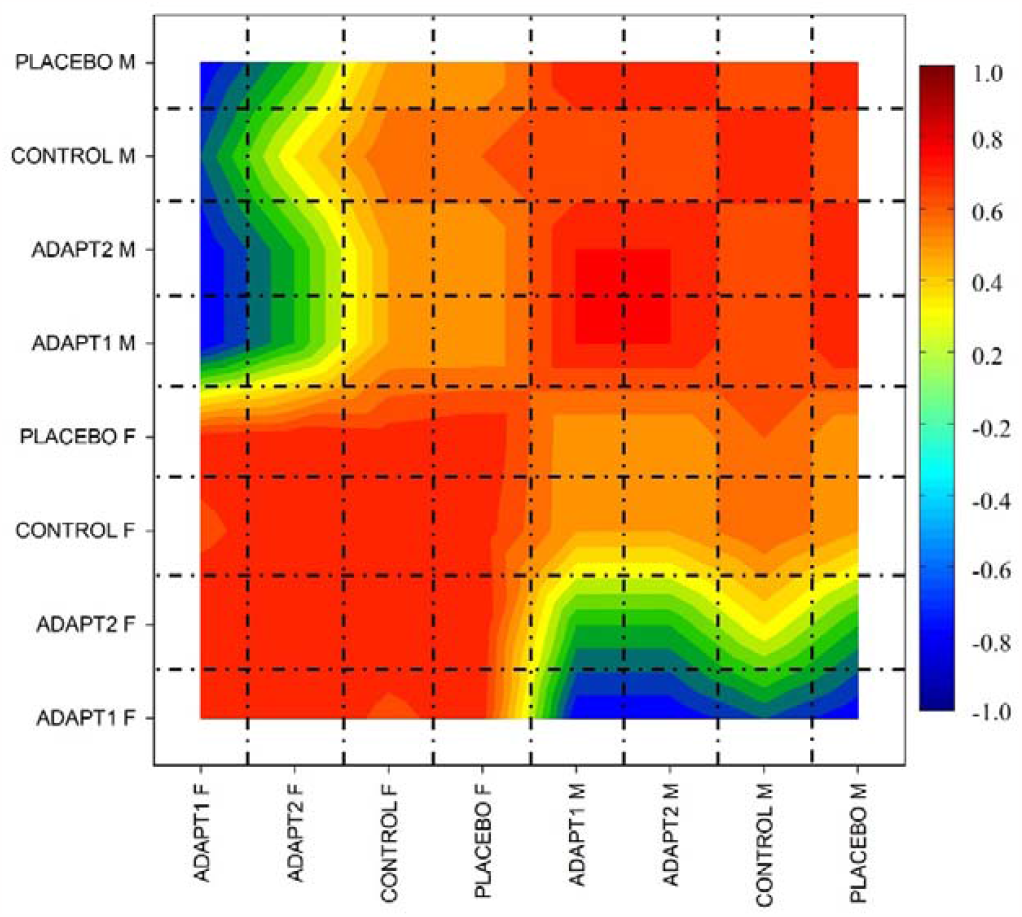
Spearman’s rank correlation test confirmed the choice of separating the men effect from women.

Not only did the spearman test confirm the separation by sex, but also the Euclidean Similarity index and the Fruchterman-Reingold Algorithm (Fig 5). The figure above shows the groupings between genders so that only when women were subjected to the placebo effect did their performance approach that of men, and women could be more easily affected by noetics.

**Figure 5.**
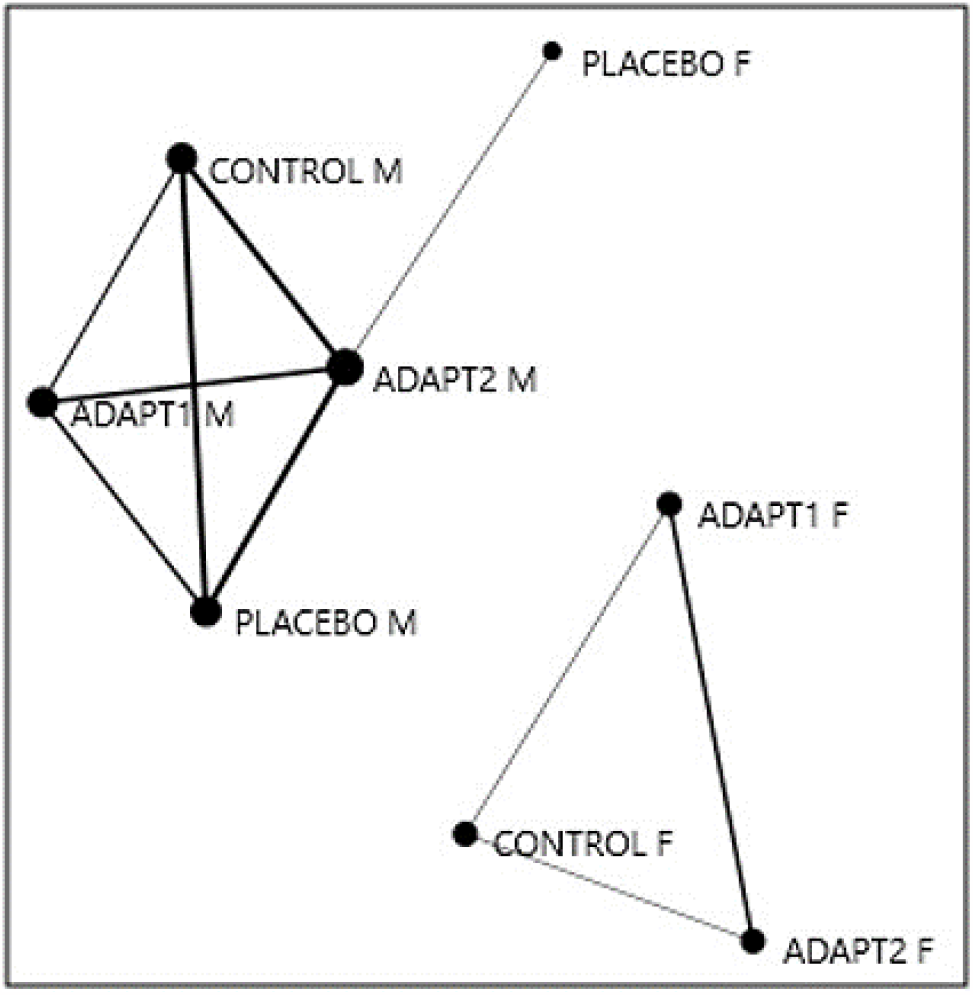
Not only did the spearman test confirm the effect separation by sex, but also the Euclidean Similarity index and the Fruchterman-Reingold Algorithm.

Finally, the adaptation in a test and the control and placebo effects are different between the sexes, from the Euclidean distance Neighbor Joining clustering (Fig 6A) and correspondence analysis (Fig 6B).

**Figure 6.**
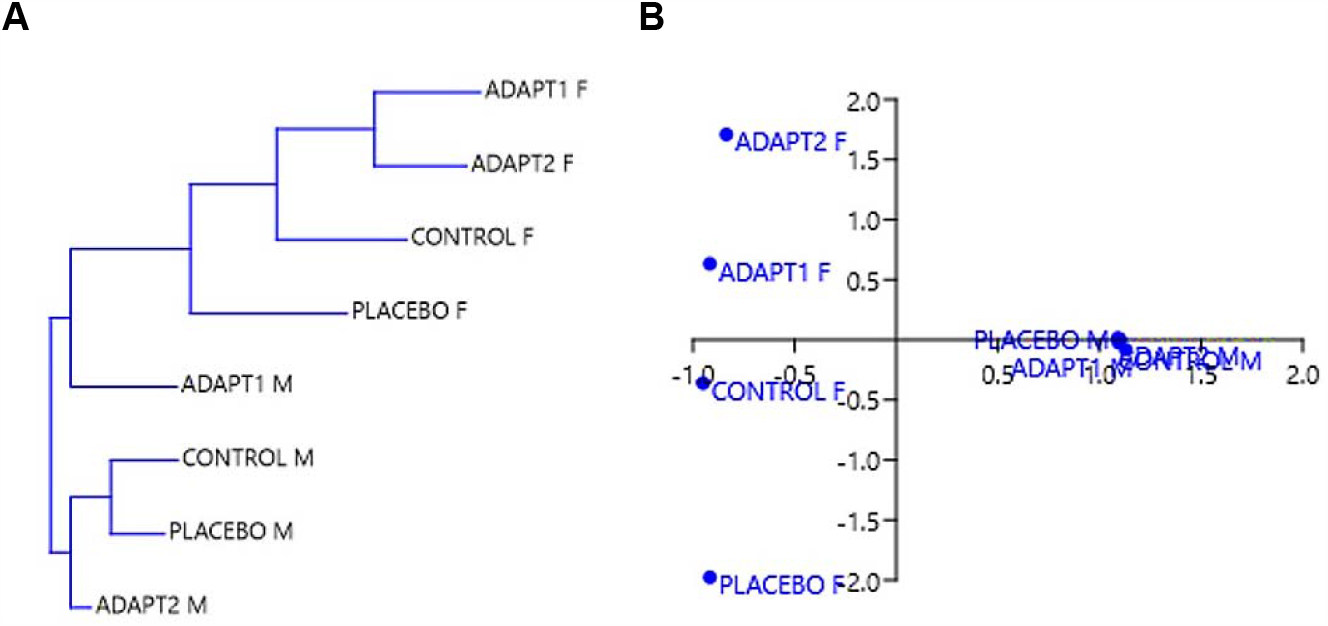
The adaptation in a test and the control and placebo effects are different between the sexes, from the Euclidean distance Neighbor Joining clustering (6A) and correspondence analysis (6B).

## Discussion

Cluster analysis was applied to group the observations into groups that are homogeneous among themselves and heterogeneous among groups through an agglomerative hierarchical method, where the number of clusters was defined throughout the analysis, and the scheme used was the nearest neighbor with a single linkage.

Regarding the unsupervised and exploratory model of machine learning, a principal component analysis (PCA) was used. Initially, Pearson’s correlation coefficient for PCA feasibility is calculated, correspondence analysis (CA) is applied, a valuable data science visualization technique to discover and display the relationship between the created categories.

Noetic Science is an area of study focused on analyzing metaphysical phenomena related to the subjectivity of human consciousness (Pekala, 2015). This awareness ranges from primarily affective forms to more advanced cognitive forms of neural processing, ranging from anoetic (non-awareness) consciousness based on affective feelings to noetic (knowledge-based) and autonoetic (higher reflective mental) processes that allow for conscious awareness (Vandekerckhove & Panksepp, 2011).

Although it is still a science on the rise, it is possible to correlate its object of study with significant advances in several areas of knowledge (Pekala, 2016), including the health area, where studies already point to its contribution as complementary medicine through the insertion of holistic practices., providing comprehensive care and thus ensuring greater efficiency in the therapeutic process (Sawni & Breuner, 2017).

Since the release of the book “The Lost Symbol” by author Dan Brown, Noetic Science has gained worldwide repercussion, so that the Institute of Noetic Sciences (IONS), located in California (USA) and founded in 1973, received an increase of the order 1000% of hits to its official website (Horrigan, 2010).

Health is a “state of complete physical, mental and social well-being and not merely the absence of disease or infirmity. This premise resumes dimensions once harmed by Cartesian dualism, as it is closely linked to the understanding of individuals as integral beings, that is, endowed with body and mind (Mehta, 2011).

Traditionally, medical care is based on drug administration to obtain results. However, drugs cannot reach the root of suffering, requiring more intimate integrative health therapy interventions that can reach points whose conventional allopathic approach is incapable, such as the spiritual, energetic, and other spheres (Steinhorn et al., 2017).

Phenomenological scope, on the part of some physicians, since they insist on perpetuating retrograde thoughts and even skeptics about the non-biological origins of pathologies.

There are sufficiently robust data on the influence of Noetic Science in the treatment of different organs and systems, such as in patients with coronary syndromes (Krucoff et al., 2001), to treat anxiety and depression (Wahbeh & Nelson, 2019), to increase survival in patients with cancer (Brunelli et al., 2012),control cortisol levels (Yount et al., 2019), alleviate pain in carpal tunnel syndrome (Yount et al., 2021), interfere with human reproduction (Richardson & Webber, 1971), improve the mental health conditions of the population during pandemics such as COVID-19 (Ojeahere et al., 2020; Gonçalves & Neto 2020) and improve immunity through the production of Natural Killer (NK) cells (Neto et al., 2021).

A physiological or psychological change can characterize the placebo effect due to the belief deposited in a treatment, healing method, or supplement for physical or cognitive improvement. This response manifests as a potential bioindicator of mental action in maintaining health (Colloca & Benedetti, 2005; Benedetti, 2013; Belcher et al., 2018; Colloca, 2019).

When the most skeptical researchers say they do not believe in Noetic Science but use a placebo in their research, they generate an apparent contradiction.

The present study shows that men might be up to eight times more affected by the control effect and three times more by the placebo effect than women.

Regarding performance, there was an antagonistic behavior concerning gender for the control effect and an agonistic effect for the placebo effect, but with less impact on women. Men also showed a faster adaptation to the test.

## Conclusion

Noetic science, always considered but never assumed by researchers, is confirmed when the present study reveals that men are more affected by the control effect and the placebo effect than women, with antagonistic behavior concerning gender for the control effect and an agonist effect for the placebo effect, but with less impact on women about performance.

### Limitations

Because a convenience sample was chosen, and after applying the inclusion and exclusion criteria, there was a reduction in the sample size. However, the results are highly relevant with the advent of selected statistical methods.

### Practical applications

Studies that use a placebo to evaluate medications, and food supplements, for different procedures or objectives must consider the noetic difference between genders to avoid wrong planning, application, and evaluation of results.

## Competing interests

The authors declare that they have no competing interests.

### Authors’ contributions

RLN, JKL, MVAV, LCOG and AMMN: essential contributions to the conception and design of the study protocol; acquisition, analysis and interpretation of data; and involvement in drafting of the manuscript. RLN, LCOG and AMMN: critical revisions for important intellectual content. All authors read and approved the final manuscript.

## Acknowledgements

We thank all participants.

## Data availability

All data generated or analyzed during this study are included in this published article.

## REFERENCES

Belcher AM, Ferré S, Martinez PE, Colloca L (2018) Role of placebo effects in pain and neuropsychiatric disorders. Prog Neuropsychopharmacol Biol Psychiatry, 87:298–306 https://doi.org/10.1016/j.pnpbp.2017.06.003

Benedetti F (2013) Placebo and the new physiology of the doctor-patient relationship. Physiol Rev, 93(3):1207–1246 https://dx.doi.org/10.1152%2Fphysrev.00043.2012

Brunelli C, Bianchi E, Murru L, Monformoso P, Bosisio M, Gangeri L, Miccinesi G, Scrignaro M, Ripamonti C, Borreani C (2012) Italian validation of the Purpose In Life (PIL) test and the Seeking Of Noetic Goals (SONG) test in a population of cancer patients. Support Care Cancer, 20(11):2775–2783 https://doi.org/10.1007/s00520-012-1399-6

Buchfuhrer MJ, Hansen JE, Robinson TE, Sue DY, Wasserman K, Whipp BB. (1983) Optimizing the exercise protocol for cardiopulmonary assessment. J Appl Physiol, 55(5):1558–1564. https://doi.org/10.1152/jappl.1983.55.5.1558

Cohen J. (1992) Quantitative methods in psychology. A power primer. Psychological Bulletin, 112(1):155–159. https://doi.org/10.1037//0033-2909.112.1.155

Colloca L, Benedetti F (2005) Placebos and painkillers: is mind as real as matter? Nat Rev Neurosci, 6(7):545–552 https://doi.org/10.1038/nrn1705

Colloca L (2019) The Placebo Effect in Pain Therapies. Annu Rev Pharmacol Toxicol, 59:191–211 https://doi.org/10.1146/annurev-pharmtox-010818-021542

Gonçalves LCO, Bessa A, Freitas-Dias R, Luzes R, Werneck-de-Castro JPS, Bassini A, Cameron LC. (2012) A sportomics strategy to analyze the ability of arginine to modulate both ammonia and lymphocyte levels in blood after high-intensity exercise. J Int Soc Sports Nutr, 9(1):30. https://doi.org/10.1186/1550-2783-9-30

Gonçalves LCO, Neto AMM (2020) The use of existing therapeutic agents to combat COVID-19. South American Journal of Basic Education, Technical and Technological, 7(2):912–921

Gonçalves LCO, Neto AMM, Bassini A, Prado ES, Muniz-Santos R, Verli MVA, Jurisica L, Lopes JSS, Jurisica I, Andrade CMB, Cameron LC. (2022) Sportomics suggests that albuminuria is a sensitive biomarker of hydration in cross combat. Scientific Reports, 12(8150):1–12. https://doi.org/10.1038/s41598-022-12079-7

Horrigan BJ (2010) The Lost Symbol sparks nationwide interest in the noetic sciences. Explore, 6(1):11–14 https://doi.org/10.1016/j.explore.2009.11.007

Krucoff MW, Crater SW, Green CL, Maas AC, Seskevich JE, Lane JD, Loeffler KA, Morris K, Bashore TM, Koenig HG (2001) Integrative noetic therapies as adjuncts to percutaneous intervention during unstable coronary syndromes: Monitoring and Actualization of Noetic Training (MANTRA) feasibility pilot. Am Heart J, 142(5):760–769 https://doi.org/10.1067/mhj.2001.119138

Mehta N (2011) Mind-body Dualism: A critique from a Health Perspective. Mens Sana Monogr, 9(1):202–209 https://doi.org/10.4103/0973-1229.77436

Neto AMM, Verli MVA, Freitas AMWH, Silva EP, Jesus GS, Silva TS, Paulino EFR, Gonçalves LCO (2021) A narrative review of the triad about stress, NK cells, and D-dimer. Brazilian Journal of Development, 7(1):2228–2239 https://doi.org/10.34117/bjdv7n1-152

Ojeahere MI, Filippis R, Ransing R, Karaliuniene R, Ullah I,Bytyçi DG, Abbass J, Kilic O, Nahidi M, Hayatudeen N, Nagendrappa S, Shoib S, Chonnakarn J, Larnaout A, Maiti T, Ogunnubi OP, Hayek SE, Bizri M, Teixeira ALS, Pereira-Sanchez V, Costa MP (2020) Management of psychiatric conditions and delirium during the COVID-19 pandemic across continents: lessons learned and recommendations. Brain Behav Immun Health, 9:1–10 https://doi.org/10.1016/j.bbih.2020.100147

Pekala RJ. (2015) Hypnosis as a “state of consciousness”: how quantifying the mind can help us better understand hypnosis. Am J Clin Hypn, 57(4):402–424 https://doi.org/10.1080/00029157.2015.1011480

Pekala RJ. (2016) The “Mysteries of Hypnosis:” Helping us better understand hypnosis and empathic involvement theory (EIT). Am J Clin Hypn, 58(3);274–285 https://doi.org/10.1080/00029157.2015.1101679

Richardson AC, Webber LD (1971) The noetic dimension of human reproduction. Am J Obstet Gynecol, 110(6):808–824 https://doi.org/10.1016/0002-9378(71)90579-5

Shapiro, AK. (1960) A contribution to a history of the placebo effect. Behavioral Science, 5:109–135. https://doi.org/10.1002/bs.3830050202

Simon R. (2000) Are placebo-controlled trials ethical or needed when alternative treatment exists? Ann Intern Med, 133(6):474–475. https://doi.org/10.7326/0003-4819-133-6-200009190-00017

Sawni A, Breuner CC. (2017) Clinical Hypnosis, an Effective Mind-Body Modality for Adolescents with Behavioral and Physical Complaints. Children (Basel), 4(4):19 https://doi.org/10.3390/children4040019

Steinhorn DM, Din J, Johnson A (2017) Healing, spirituality and integrative medicine. Ann Palliat Med, 6(3):237–247 https://doi.org/10.21037/apm.2017.05.01

Turner M (2015) Can the Effects of Religion and Spirituality on Both Physical and Mental Health be Scientifically Measured? An Overview of the Key Sources, with Particular Reference to the Teachings of Said Nursi. J Relig Health, 54(6):2045–2051 https://doi.org/10.1007/s10943-014-9894-3

Vandekerckhove M, Panksepp J. (2011) A neurocognitive theory of higher mental emergence: from anoetic affective experiences to noetic knowledge and autonoetic awareness. Neurosci Biobehav Rev, 35(9):2017–2025 https://doi.org/10.1016/j.neubiorev.2011.04.001

Wahbeh H, Nelson M (2019) iRest Meditation for Older Adults with Depression Symptoms: A Pilot Study. Int J Yoga Therap, 29(1):9–17 https://doi.org/10.17761/2019-00036

Yount G, Church D, Rachlin K, Blickheuser K, Cardonna I (2019) Do Noncoding RNAs Mediate the Efficacy of Energy Psychology? Glob Adv Health Med, 8:1–8 https://dx.doi.org/10.1177%2F2164956119832500

Yount G, Delorme A, Radin D, Carpenter L, Rachlin K, Anastasia J, Pierson M, Steele S, Mandell H, Chagnon A, Wahbeh H (2021) Energy Medicine treatments for hand and wrist pain: A pilot study. Explore, 17(1):11–21 https://doi.org/10.1016/j.explore.2020.10.015

